# Cortico-subcortical β burst dynamics underlying movement cancellation in humans

**DOI:** 10.1101/2021.02.06.430074

**Authors:** Darcy A. Diesburg, Jeremy D. W. Greenlee, Jan R. Wessel

**Affiliations:** Department of Psychological and Brain Sciences, University of Iowa, Iowa City, IA, USA; Department of Neurosurgery, University of Iowa Carver College of Medicine, Iowa City, IA, USA; Iowa Neuroscience Institute, University of Iowa, Iowa City, IA, USA; Department of Neurology, University of Iowa Carver College of Medicine, Iowa City, IA, USA

**Keywords:** movement cancellation, inhibitory control, β bursts, STN, thalamus

## Abstract

Dominant neuroanatomical models hold that humans regulate their movements via loop-like cortico-subcortical networks, including the subthalamic nucleus (STN), thalamus, and sensorimotor cortices (SMC). Inhibitory commands across these networks are purportedly sent via transient, burst-like signals in the β frequency (15-29Hz). However, since human depth-recording studies are typically limited to one recording site, direct evidence for this proposition is hitherto lacking. Here, we present simultaneous multi-site depth-recordings from SMC and either STN or thalamus in humans performing the stop-signal task. In line with their purported function as inhibitory signals, subcortical β-bursts were increased on successful stop-trials and were followed within 50ms by increased β-bursting over SMC. Moreover, between-site comparisons (including in a patient with simultaneous recordings from all three sites) confirmed that β-bursts in STN precede thalamic β-bursts. This provides first empirical evidence for the role of β-bursts in conveying inhibitory commands along long-proposed cortico-subcortical networks underlying movement regulation in humans.

## Introduction

Movement cancellation – i.e., the ability to stop ongoing movements when necessary – allows humans to adapt their behavior quickly in response to changing environmental demands. This ability, which is underpinned by inhibitory control processes that operate on active motor representations, allows for flexible pursuit of short- and long-term goals. It also is often critically important in immediately preventing bodily harm, such as when one stops walking when a “don’t walk” signal flashes at a busy crosswalk.

The pathways underlying movement cancellation in the brain comprise a highly specific, well-characterized fronto-basal ganglia network for motor and cognitive control (Wessel and Aron, 2017). A predominant paradigm used by neuroscientists to probe this network and investigate inhibitory control is the stop-signal task (SST), wherein participants are tasked with making responses that they sometimes have to attempt to cancel after being presented with a sudden “stop signal” (Logan and Cowan, 1984; Verbruggen et al., 2019). Horse-race models of the SST allow for the computation of the duration of the latent cancellation process (stop-signal reaction time, SSRT), even though no overt response is observable when participants successfully stop (Verbruggen and Logan, 2008; Boucher et al., 2007.) When a stop-signal occurs, the right inferior frontal cortex (rIFC) purportedly excites the subthalamic nucleus (STN) via a monosynaptic white matter tract called the hyperdirect pathway (Nambu, Tokuno, and Takada, 2002; Aron et al., 2007; Chen et al., 2020). Subsequently, the STN broadly and non-selectively excites the internal segment of the globus pallidus (GPi), the output nucleus of the basal ganglia. In turn, the GPi then broadly inhibits the motor thalamus. It has been proposed that the resultant net inhibition of thalamocortical signaling loops (i.e., motoric loops between thalamus and sensorimotor cortex) – and the subsequent degradation of motor programs held in those loops – enables the type of rapid movement cancellation found in tasks like SST (Parent and Hazrati, 1993; Jahanshahi et al., 2015).

Electrophysiological recordings from nodes of this fronto-basal ganglia network have revealed that network communication through these pathways likely occurs in the β frequency band. During movement execution, decreases in average β power are observed over sensorimotor cortex (SMC; both intracranially and on the scalp, Crone et al., 1998; Pfurtscheller and Lopes da Silva, 1999; Kühn et al., 2004; Takemi et al., 2013) and in subcortical motor regions such as the STN (Alegre et al., 2005). In contrast, average β power is *increased* in SMC and STN when inhibitory control is required, both following stop-signals in the SST (Wessel et al., 2016; Swann et al., 2009, 2011; Ray et al., 2012; Alegre et al., 2013; Benis et al., 2014; Bastin et al., 2014) and during motor conflict more broadly (Brittain et al., 2012; Wessel, Waller, and Greenlee, 2019). Similar increases in β power during movement cancellation can be observed in cortical regions that are ostensibly upstream of the STN and thalamus in the purported cortico-subcortical loop underlying rapid movement cancellation, such as the pre-supplementary motor area (Swann et al., 2012; Picazio et al., 2014) and the rIFC (Swann et al., 2009). Together, these findings have established cortical and subthalamic β activity as an index of inhibitory control.

However, within the past several years, cross-species research has indicated that these changes in β power do not reflect *sustained* β oscillations at the single-trial level (Feingold et al., 2015; Sherman et al., 2016; Shin et al., 2017; Tinkhauser et al., 2018; Maling et al., 2018; Cagnan et al., 2019). Instead, single-trial variations in β activity are highly transient, lasting approximately 100-150ms on average – that is, unaveraged β activity has properties that is better characterized as intermittent bursting instead of slow-and-steady modulations of amplitude (van Ede et al., 2018). In line with this, β bursts are more predictive of behavior than average fluctuations in β power. For example, perceptual stimuli preceded closely by β bursts in somatosensory cortex are less likely to be detected (Shin et al., 2017) and β bursts in motor cortex closely preceding imperative stimuli are associated with slower responses (Little et al., 2019). Computational models built with biophysical principles in mind suggest that these bursts in SMC relate specifically to proximal and distal excitatory drive to the synapses of neocortical pyramidal neurons (Sherman et al., 2016). Thus, not only do β bursts carry fine-grained information about behavior on the single-trial level, they also relate more closely to underlying mechanisms than sustained β averaged over many trials. Notably, two recent studies have demonstrated that β bursts on the scalp relate specifically to the inhibitory aspects of movement regulation. One study, which included scalp EEG recordings during the SST from over 200 individuals, demonstrated reductions in β burst rates over SMC during go trials, as well as increases in burst rates over frontocentral and motor cortices during stop trials (Wessel, 2020). Furthermore, increases in burst rates over frontocentral cortex dissociated success of cancellation – in the same study, successful stop trials featured an increased number of frontocentral β bursts before SSRT on average than failed stop trials. A subsequent study by Jana and colleagues (2020) demonstrated that β bursts over prefrontal cortex were followed within 20ms by broad skeleto-motor suppression and within 40ms by outright cancellation detectable at the motor effector.

While these studies identify potential (pre)frontal cortical control signals associated with movement cancellation, by their nature as scalp-level investigations, these studies are uninformative regarding the downstream basal ganglia-thalamic dynamics through which inhibitory control of SMC is ostensibly implemented. Transient β bursts are indeed known to exist in the STN, and pathologically long β bursts are correlated with abnormal movement in individuals with Parkinson’s disease (PD; Torrecillos et al., 2018; Lofredi et al., 2019). Still, it is unclear what role subcortical β bursts play during movement regulation, and whether their subcortical dynamics conform to the dominant neurophysiological and neuroanatomical models of inhibitory control.

Our aims for the current study were twofold. Firstly, we investigated whether β bursts in subcortical regions of basal ganglia-thalamic inhibitory pathways are associated with movement cancellation. To this end, we assessed the relationship between stop-signal task performance and β burst rates in both STN and motor thalamus. We then investigated whether these subcortical bursts have reliable temporal relationships with movement-related β bursts in SMC that are suggestive of an inhibitory influence of the subcortical regions on SMC. Secondly, we evaluated existing models of inhibitory control networks by assessing relative timing of bursts from different subcortical recording sites. The dominant model of a fronto-basal ganglia circuit for inhibitory control suggests that movement cancellation is accomplished by downstream net-inhibition of the motor thalamus (Jahanshahi et al., 2015). Hence, we expected β bursts related to movement cancellation to emerge first in the STN, followed by bursts in the thalamus.

To achieve these goals, we collected recordings of local field potentials (LFPs) simultaneously from SMC and a subcortical site (either the STN or thalamus) during awake deep-brain stimulation (DBS) lead implantation surgery in two groups of patients: PD patients undergoing STN implantation and essential tremor patients undergoing implantations in the motor thalamus (i.e., ventral intermediate nucleus, VIM). Moreover, data from one highly unique PD patient included simultaneous recordings from all three locations: SMC, STN, and VIM. During the recordings, patients performed an auditory version of the stop-signal task to test their ability to rapidly cancel movements. This study is the first in humans to investigate the dynamics of cortical and subcortical β bursts – as well as their multi-site interactions – during movement cancellation.

## Results

### Behavior

While local field potentials were recorded from SMC and either STN or VIM, 21 participants completed an auditory version of the SST (**see Figure 1**). Behavioral results can be found in **Table 1**. Go accuracy was the only behavioral metric which significantly differed between the STN and VIM DBS groups (*T*(19) = 5.22, *p* < .0001, *d* = 2.27), with VIM patients responding more accurately on go-trials. On average, the VIM implant group also responded faster during correct go and failed stop trials and cancelled movements more quickly than STN DBS patients, as indicated by SSRT (which was in the typical elongated range for movement disorder patients: Gauggel et al., 2004; Obeso et al., 2011; Hughes et al., 2019). However, none of these results were significant, indicating comparable task performance between both groups.

**Table 1.** Means of stop-signal task behavioral performance metrics, across and within patient groups. * Indicates significant difference between VIM and STN groups at p < .0001.

|  | Reaction times (ms) |  |  | Accuracy |  |
| --- | --- | --- | --- | --- | --- |
|  | Go | Failed stop | SSRT | Go trials* | Stop trials |
| All participants | 943 | 763 | 474 | 0.89 | 0.61 |
| STN DBS | 952 | 786 | 533 | 0.83 | 0.55 |
| Thalamic DBS | 934 | 741 | 421 | 0.94 | 0.65 |

**Figure 1.**
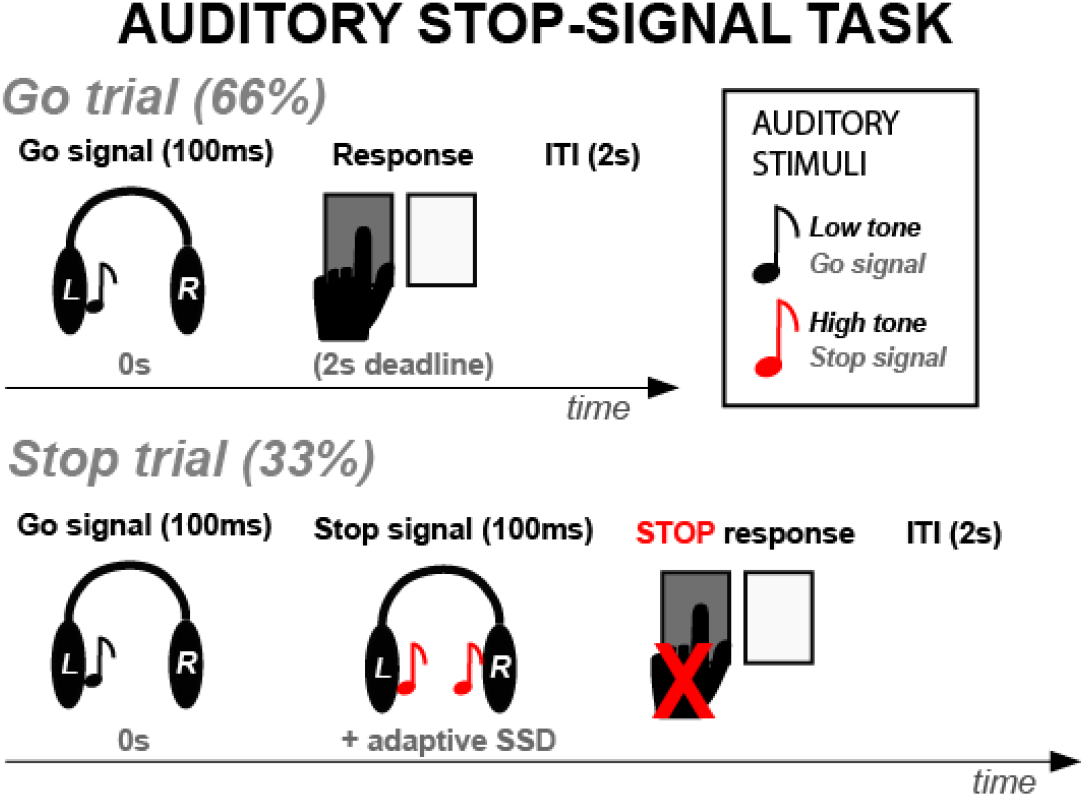
Participants performed an auditory version of the stop-signal task while LFP recordings were collected.

### ERSPs

To confirm the electrode placement over SMC, event-related spectral perturbation (ERSP) was quantified in the window ranging from −100 to 1500ms relative to the go-signal on go trials. In line with the expected pattern (cf., Crone et al., 1998; Pfurtscheller and Lopes da Silva, 1999; Kühn et al., 2004; Takemi et al., 2013), decreases in trial-averaged β band (15-29Hz) power were observed at contralateral (to the response) SMC sites following go signals (see **Figure 2**). Moreover, this pattern was evident in both the localizer task as well as the stop-signal task.

**Figure 2.**
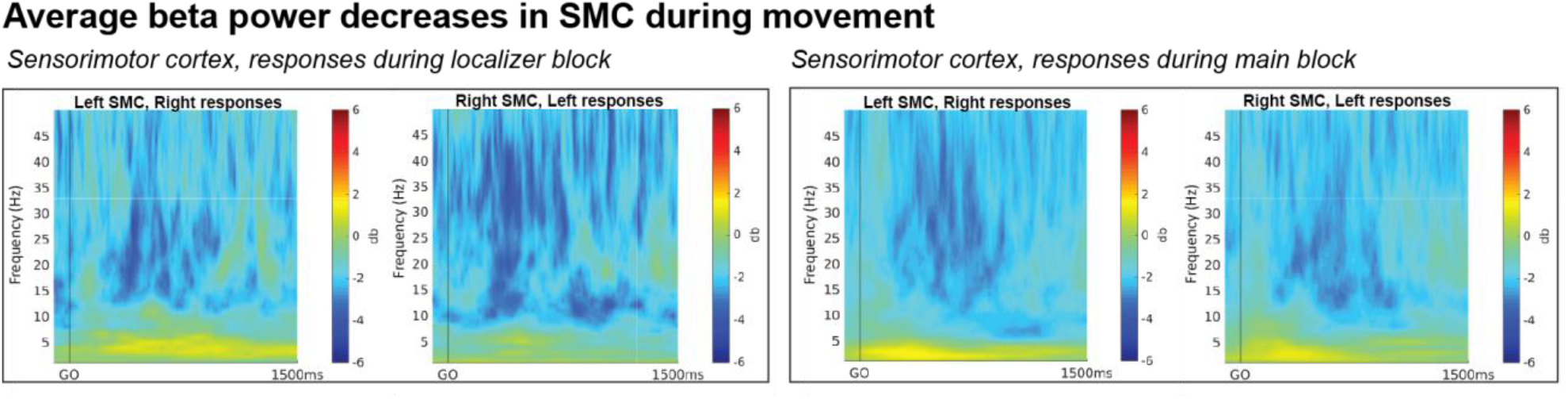
Average ERSPs during movement on go-trials, time-locked to go signals. Both sessions show clearly visible β power decreases following the go signal.

### *β* bursts decrease during movement initiation

We then investigated whether the SMC sites showed the corresponding pattern of movement-related reductions in β burst rates in the lead-up to the response (Wessel et al., 2020; Soh et al., 2021), and whether that pattern was also present in the subcortical nuclei. Burst rates were quantified in 9 non-overlapping bins of 100ms starting from the onset of the go signal. Indeed, we observed reductions in β burst rates during movement execution in all recording locations. A 3 × 2 ANOVA revealed a significant effect of *TIMEPOINT* on burst rate, observed in STN (*F*(2,8) = 8.78, *p <* .0001, *η*^*2*^ = .31), VIM (*F*(2,10) = 6.79, *p* < .0001, *η*^*2*^ = .25), and in the SMC (*F*(2,19) = 8.04, *p* < .0001, *η*^*2*^ = .22; **see Figure 3A)**.

**Figure 3.**
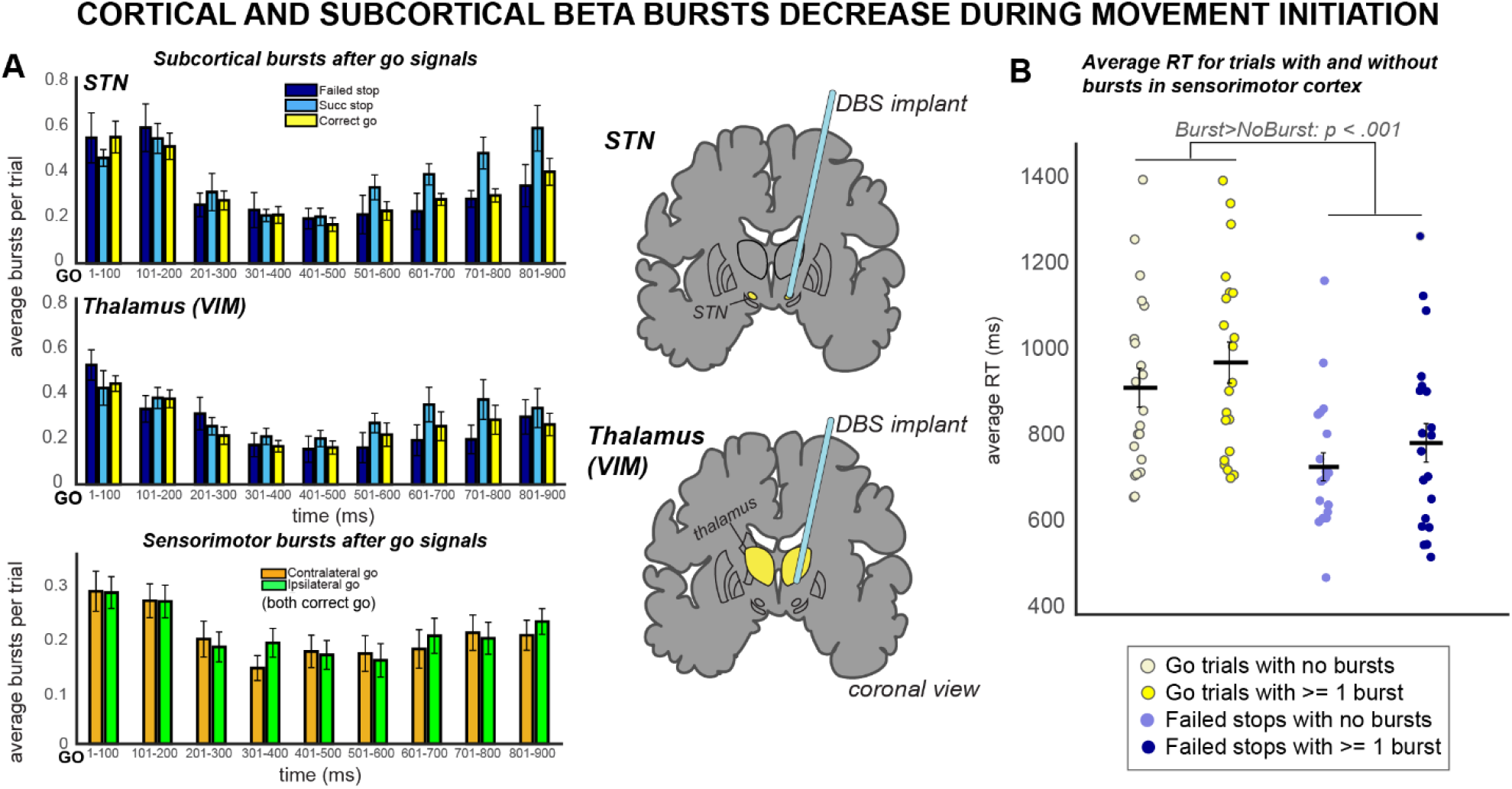
*β* burst rates decreased across recording sites during movement. A) *β* burst rates following the go signal on go-trials in the stop-signal task, displayed at all three recording sites. Burst rates decrease quickly following the go signal in STN, thalamus, and SMC. B) The occurrence of *β* bursts in SMC between the go signal and the response was associated with slower reaction times.

In the STN group, there was also a significant effect of *TRIAL TYPE* (*F*(2,8) = 4.95, *p =*.02, *η*^*2*^ = .02) on burst rate, but no *TRIAL TYPE × TIMEPOINT* interaction (*F*(2,8) = 1.58, *p =* .08, *η*^*2*^ = .05). On the other hand, in the VIM group, there was no significant effect of *TRIAL TYPE* (*F*(2,10) = 2.85, *p* = .08, *η*^*2*^ = .02) on burst rate, but there was a significant *TRIAL TYPE × TIMEPOINT* interaction (*F*(2,4) = 1.77, *p =* .04, *η*^*2*^ = .05).

These findings are in line with the proposition that β bursts are related to an inhibited state of the motor system, which must be downregulated to enable movement through a net-disinhibition of the cortico-subcortical motor circuitry (e.g., Soh et al., 2021).

### β bursts have inhibitory effects on response times

Further evidence for the assertion that β bursts relate to an inhibited state of the motor system comes from analyses that have shown that movements that are closely preceded by a β burst in SMC are slower than movements that are not preceded by a burst (Shin et al., 2017; Little et al., 2019). We repeated this analysis in our sample. Indeed, for both failed stop and correct go trials, trials containing β bursts between the go signal and the response had significantly longer mean reaction times (**see Figure 3B**). A 2 × 2 ANOVA revealed significant effects of *TRIAL TYPE* (*F*(2,19) = 40.07, *p* < .0001, *η*^*2*^ = .56) and *BURST PRESENCE* (*F*(2,19) = 19.67, *p* = .0002, *η*^*2*^ =.05) on reaction time, though a significant interaction between the two was not found (*F*(4,19) =.009, *p* = .92, *η*^*2*^ < .001). Hence, in line with prior findings, cortical β bursts in our sample appear to index a greater degree of net-inhibition in SMC prior to movement initiation.

### *β* bursts increase during movement cancellation

The abovementioned analyses confirmed that the SMC electrodes were accurately placed and that all sites displayed reasonable β burst dynamics that would be expected under the assumption that β bursts relate to an inhibited state of the motor system.

The main hypothesis of our paper was that after stop-signals in the stop-signal task, motor inhibition is achieved by a rapid re-instantiation of an inhibited state in SMC, preceded by β Burst signaling from the subcortical nuclei. The first test of this hypothesis was to investigate whether successful stop-trials were accompanied by an increase in subcortical β bursting compared to matched go-trials and failed stop-trials.

Indeed, during the critical time period between the onset of the stop signal and the end of SSRT, a significant main effect of *TRIAL TYPE* (successful stop, failed stop, matched go) on burst count was found at both subcortical sites (STN: *F*(9) = 6.83, *p* = .006, *η*^*2*^ = .43; VIM: *F*(11) = 6.52, *p* = .006, *η*^*2*^ = .37). Follow-up pairwise *t-*tests revealed that this was due to a significant increase in bursts in the stop signal delay (SSD)-SSRT period on successful stop trials compared to matched go trials (STN: *p* = .01; VIM: *p* = .01) and increased bursts on successful stop trials compared to failed stop trials (STN: *p* = .04; VIM: *p* = .03; **see Figure 4**). This is in line with the assumption that early-latency subcortical β burst signaling reflects a rapid deployment of inhibitory control after a stop-signal.

**Figure 4.**
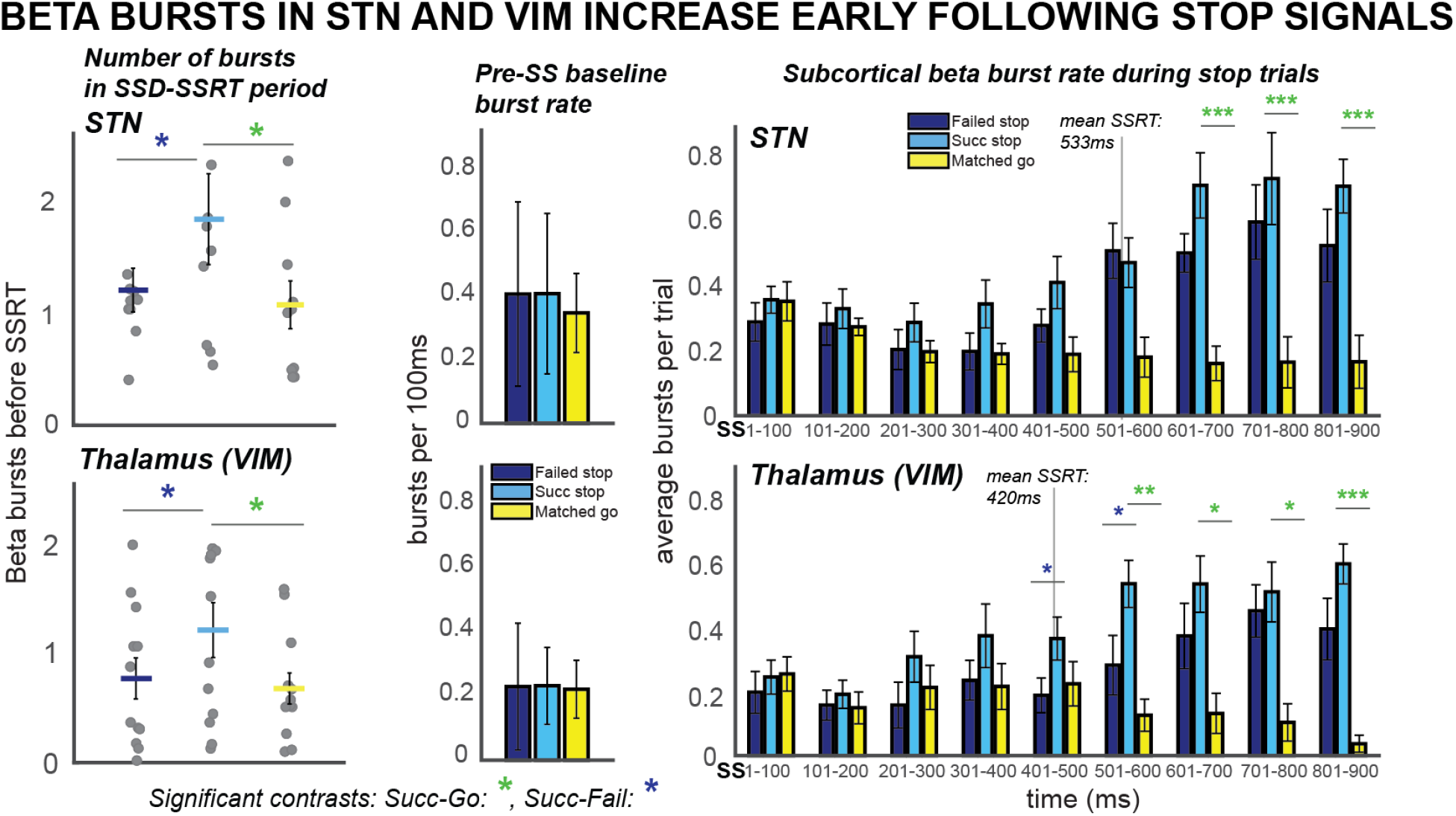
*β* burst rates increase following stop signals. Burst rates increased at early latencies in STN and VIM during successful cancellation, as well as at later latencies in both VIM and STN during stop trials. These differences could not be accounted for by differences in pre-stop baseline burst rates. The number of total bursts in the SSD-SSRT time period in STN and VIM differ across trial types, with a greater number of bursts before SSRT for successful stops compared to go trials. (* indicates p < .05, ** indicates p < .01, *** indicates p < .001.)

As a control analysis, we also compared burst rates in the baseline period before the stop signal. A 3 × 1 ANOVA revealed no significant effects of *TRIAL TYPE* on β bursts in the pre-stop baseline for either STN or thalamus (STN: *F*(2,8) = 0.25, *p =* .78, *η*^*2*^ = .03; VIM:: *F*(2,10) = 0.02, *p =* .98, *η*^*2*^ = .002). Hence, the stop-signal related differences in burst rates were not attributable to differences in the baseline.

To map out β burst dynamics in the post-go and post-stop periods in more temporal detail, we also calculated the average burst rate in successive, non-overlapping time bins of 100ms covering the 1000ms period starting 100ms before the stop-signal. While this does not take into account each participants’ SSRT, it does provide a more comprehensive picture of the development of subcortical β bursting over time. In the STN, we observed significant effects of *TRIAL TYPE* (*F*(2,8) = 35.14, *p <* .0001, *η*^*2*^ = .19) and *TIMEPOINT* (*F*(2,8) = 4.19, *p =* .004, *η*^*2*^ =.13), as well as a significant *TRIAL TYPE × TIMEPOINT* interaction (*F*(4,8) *=* 5.87, *p* < .0001, *η*^*2*^= .14). The same pattern was observed in the VIM, again with significant main effects of *TRIAL TYPE* (*F*(2,10) = 26.97, *p <* .0001, *η*^*2*^ = .20) and *TIMEPOINT* (*F*(2,10) = 3.16, *p =* .004, *η*^*2*^ = .08), and a *TRIAL TYPE × TIMEPOINT* interaction (*F*(4,10) *=* 6.00, *p* < .0001, *η*^*2*^ = .15).

Pairwise follow-up *t-*tests were then used to probe differences between successful stops and go trials and between successful and failed stop trials at individual time bins. Burst rates for successful stop trials were significantly greater than for go trials at 601-700ms (*p* < .001), 701-800ms (*p* < .001), and 801-900ms (*p* < .001) following SSD in the STN and at 501-600ms (*p* =.002), 601-700ms (*p* = .02), 701-800ms (*p* = .02), and 801-900ms (*p* < .001) following SSD in the thalamus (**see Figure 4**). In the VIM, there were also significant differences between successful and failed stop trial burst rates at 401-500 (*p* = .04) and 501-600ms (*p* = .05) following SSD. Note that there was no increase in early-trial β bursting on successful stop-trials in this bin-wise quantification, which is at odds with the above-mentioned quantification that measured β burst counts in each individual subjects’ stop-signal-to-SSRT period. This again suggests a tight relationship between β burst dynamics and inhibitory control behavior – resulting in the fact that differences in β burst rates after stop-signals are obscured when between-subject differences in SSRT are not taken into account.

### STN β bursts upregulate SMC bursts during cancellation

The proposition that inhibitory control in the SST is implemented via a rapid re-instantiation of SMC inhibition caused by bursts in STN/Thalamus implies that SMC β burst rates should be increased in the immediate aftermath of subcortical bursts (cf., Wessel, 2020 for a demonstration of the same relationship between fronto-central scalp-recorded β bursts and subsequent β-burst rates over SMC). To test this, we quantified SMC β bursting time-locked to the occurrence of any subcortical β bursts that occurred in the first 500ms following the stop-signal. Indeed, there was a main effect of *TIMEPOINT* (*F*(2,8) = 5.22, *p* = .006, *η*^*2*^ = .14) on SMC bursts time-locked to STN bursts. Moreover, follow-up *t*-tests revealed that bursts in STN were followed within 50ms by a difference between burst rates for successful stops compared to go trials, with burst rates increasing for successful stops, though this difference did not survive multiple-comparisons corrections (uncorrected *p* = .03; see **Figure 5**).

**Figure 5.**
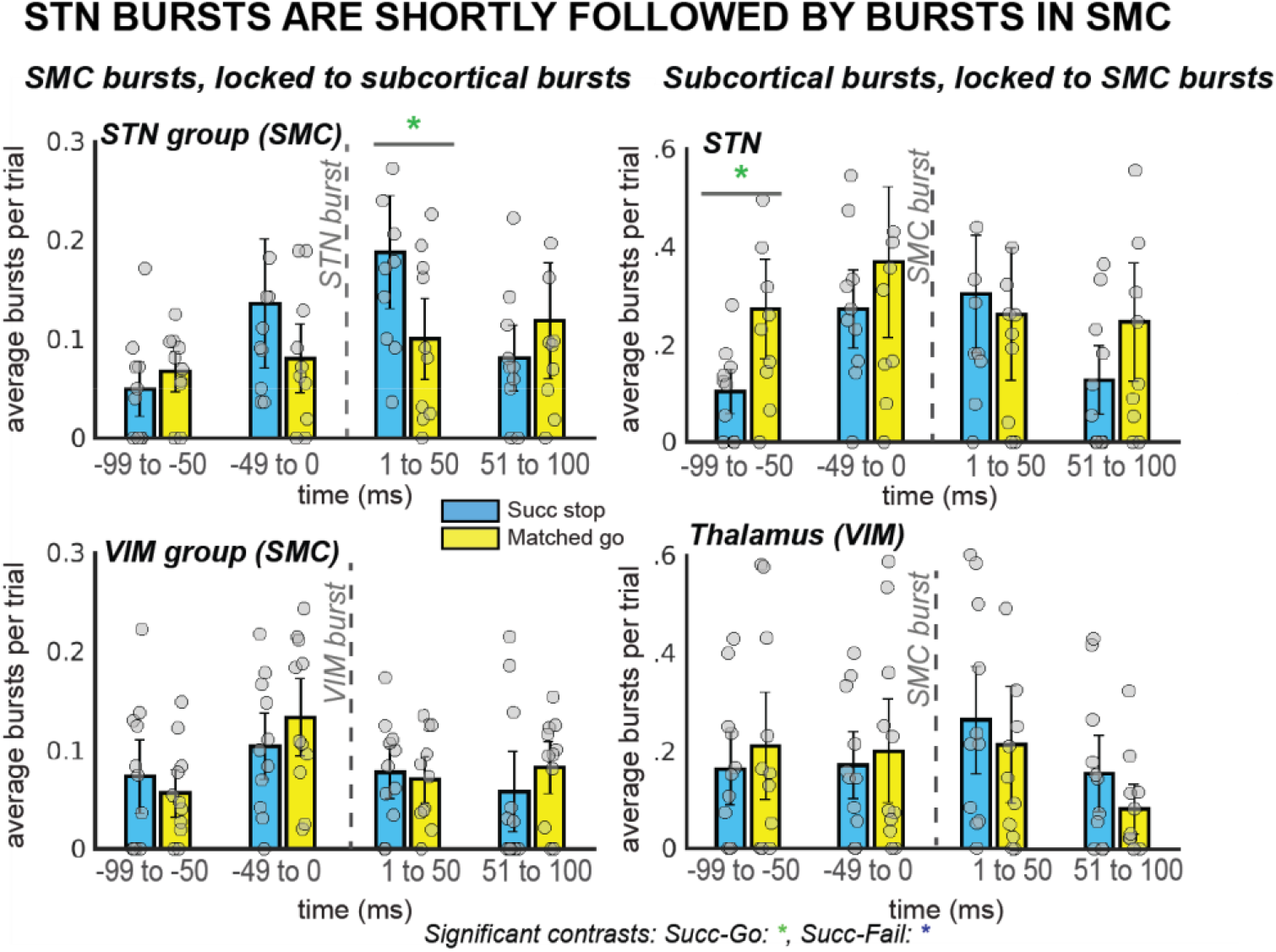
*β* burst rates in SMC increase following STN bursts. In the STN group, the first STN burst following the stop signal (or SSD, for matched-go trials) was followed within 50ms by bursts in the SMC for stop, but not go, trials. This reliable temporal relationship between STN and SMC bursts during movement cancellation did not follow the opposite pattern – STN bursts did not reliably follow SMC bursts at a specific time point. (* indicates uncorrected p < .05)

Our 2 × 2 ANOVA did not reveal a significant effect of *TRIAL TYPE* (*F*(2,8) = 0.54, *p* =.59, *η*^*2*^ = .01) or a *TRIAL TYPE × TIMEPOINT* interaction (*F*(4,8) = 1.94, *p* = .09, *η*^*2*^ = .07) for SMC bursts time-locked to STN bursts. Likewise, no effects of *TRIAL TYPE* (*F*(2,10) = 1.55, *p* = .24, *η*^*2*^ = .05), *TIMEPOINT* (*F*(2,10) = 1.71, *p* = .19, *η*^*2*^ = .03), or an interaction (*F*(4,10) = 0.81, *p* = .58, *η*^*2*^ = .04) were found for SMC bursts time-locked to VIM bursts. There were no significant pairwise differences between burst rates in VIM for successful stops versus failed go trials.

Conversely to the increase in SMC burst rate after STN bursts, we did not see an increase of STN or VIM β bursts *following* SMC bursts. A 2 × 2 ANOVA of burst rates in STN and VIM time-locked to SMC bursts revealed an effect of *TIMEPOINT* in both regions (STN: *F*(2,8) = 3.73, p = .02, *η*^*2*^ = .08; VIM: *F*(2,10) = 3.10, p = .04, *η*^*2*^ = .05) and an effect of TRIAL TYPE in VIM (VIM: *F*(2,10) = 5.32, p = .01, *η*^*2*^ = .13), but no significant main effects of *TRIAL TYPE* in STN (*F*(2,8) = 0.76, p = .48, *η*^*2*^ = .03) or interactions between the two factors (STN: *F*(4,8) = 0.69, p = .66, *η*^*2*^ = .03; VIM: *F*(4,10) = 1.12, p = .36, *η*^*2*^ = .04). No significant pairwise comparisons were found for burst rates during successful stops compared to failed stops at individual time points in either region, though a difference at the -100 to -50ms timepoint in the STN was significant before multiple comparisons correction (uncorrected *p* = .02).

The observation of elevated SMC β bursts following, but not preceding, STN bursts supports for the proposition that subcortical bursts lead to a rapid upregulation of SMC bursts during stopping.

### STN β bursts precede VIM bursts during cancellation

A key prediction of existing network models of inhibitory control is that STN is upstream from VIM – i.e., during the purported cascade that results in movement cancellation, STN signaling should temporally precede VIM signals. To test whether this is the case for the β burst signals observed here, we calculated the average latency of the first β burst after the stop signal for each subcortical recording site and compared them between groups, as well as within a single subject with implants at both sites. To account for differences in SSRT across participants, we quantified the onset latency of first bursts with respect to participant-wise SSRT. Across the group-level sample, STN bursts on average occurred before VIM bursts during failed and successful stop trials (see **Figure 6A**). While there was a significant effect of first burst time on TRIAL TYPE (with the first burst occurring earlier on successful stop trials compared to failed stop-rials in both VIM and STN; *F(1,19)* = 8.33; *p* = 01; *η*^*2*^ = .02), there was no significant effect of *LOCATION* on average burst timing (*F(1,19)* = 1.90; *p* = .18; *η*^*2*^ = .08), and no interaction (*F(1,19)* = 0.42; *p* = .52; *η*^*2*^ = .001).

**Figure 6.**
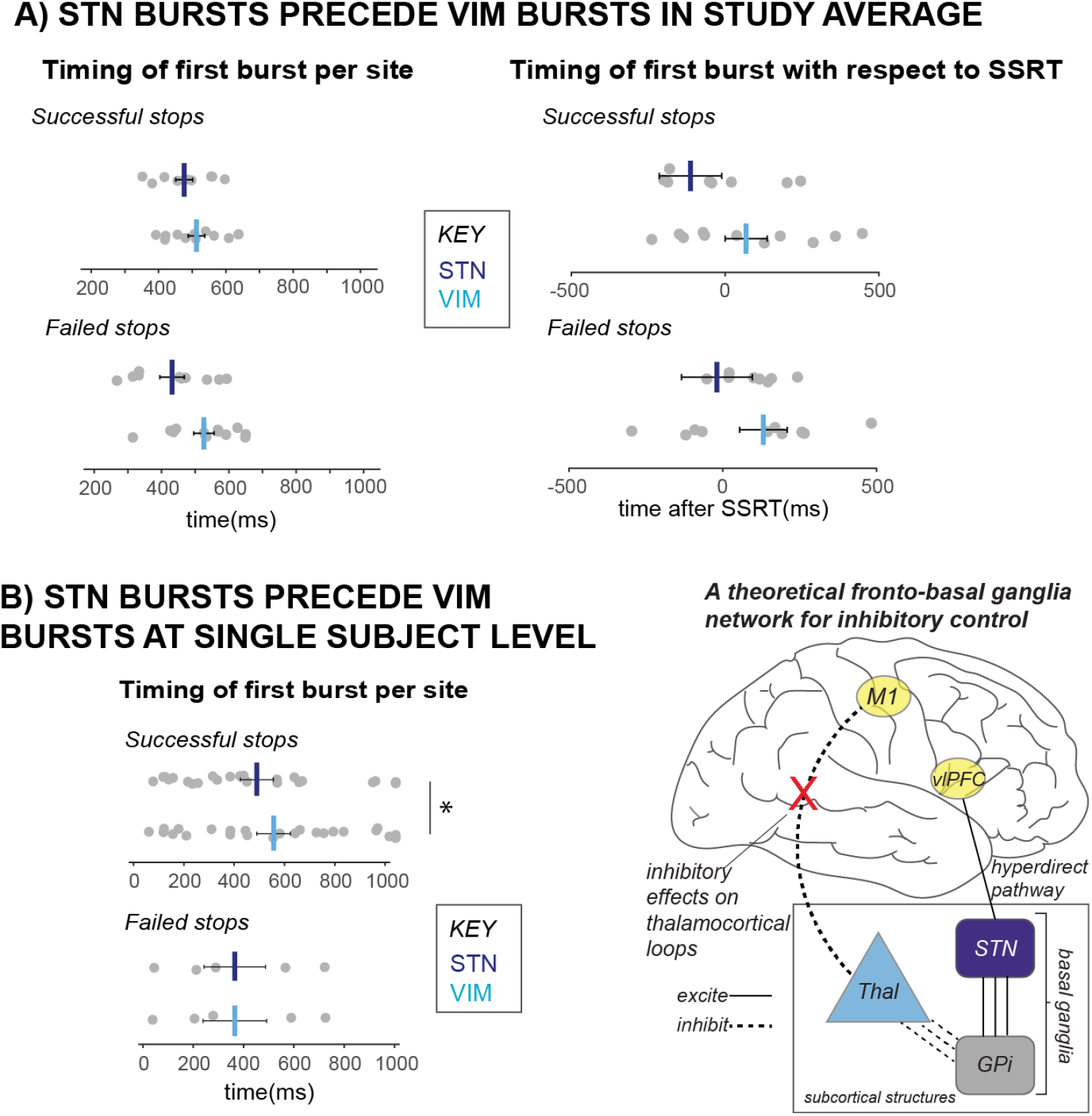
*β* bursts in STN precede bursts in VIM. Across the entire study sample (a) and in our single subject with both STN and VIM recording sites (b), STN bursts occurred earlier than VIM bursts during cancellation. These findings from subcortical regions in our datasets lend support for an account of subcortical dynamics proposed in a theorized network model of movement cancellation, which posits that the STN is recruited prior to and acts to net-inhibit the thalamus during cancellation.

However, the ultimate test of burst timing differences across regions is provided by the single subject who had recordings from both regions, as this provides the only comparison with equal behavior (specifically, SSRT). In line with the qualitative pattern observed in the group-level comparison, in this single subject, bursts in the STN occurred on average before bursts in the VIM during stop trials (see **Figure 6B**), with pairwise *t*-tests revealing significant differences between burst timing in STN and VIM during successful stop trials (*p* = .03), but not during failed stop trials (*p* = .21). This observation that cancellation-related STN bursts occur before bursts in the thalamus also supports accounts that movement regulation may be accomplished by STN-facilitated inhibition of the thalamus during a period before behavioral cancellation is observed (i.e., before SSRT).

## Discussion

We used simultaneous, multi-site intracranial recordings in awake, behaving humans to delineate the cortico-subcortical β-burst dynamics that underlie the inhibitory control of movement. Our findings have significant implications for our understanding of inhibitory control in the human brain, as well as the nature of β signaling in human motor circuitry.

### *β* bursts in the STN and thalamus relate to movement cancellation

β bursts in the human STN and thalamus relate to the rapid deployment of inhibitory control during movement cancellation in the stop-signal task. An analysis of subcortical β burst counts between the stop signal and SSRT revealed a greater number of bursts present during successful stops than during failed stops or matched go trials. Moreover, the analysis of first-burst timing revealed that in both STN and thalamus, the timing of the first burst distinguished successful from failed stopping, as earlier bursts lead to significantly higher rates of stopping. Our findings in intraoperative movement disorder participants mirror results recently obtained from scalp recordings in healthy subjects (Wessel, 2020, Jana et al., 2020), which demonstrated increased and earlier β bursts during movement cancellation at the cortical sites that are ostensibly up-stream of the basal ganglia circuitry investigated here (cf., Chen et al., 2020). The current study is the first to concretely demonstrate the relationship between subcortical β bursts and movement cancellation in the human brain. Thereby, it fills a substantial gap in our knowledge regarding the chain of neurophysiological processes in purportedly inhibitory cortico-basal ganglia-thalamic circuits.

Notably, this finding is – superficially – at odds with a recent report from Mosher and colleagues (Mosher et al., 2021), who, amongst other things, purportedly showed that human STN and SMC β bursts appear to be dissociated from activity in movement-associated neurons. Specifically, while in their data, movement-related neurons in STN showed reduced firing prior to movement cancellation during successful stops, β bursts were not observed until later latencies. However, while the finding of later-latency β bursting in their study is replicated here, the apparent discrepancy in the early-trial β burst rates is a reflection of key differences in the way β bursts were measured with respect to the stop-signal. Specifically, Mosher and colleagues quantified β burst rates in a continuous manner after the stop signal, using sliding windows. Since SSRT varies considerably across participants (particularly in clinical samples), using such a quantification does not take into account the substantial between-subject variance in SSRT (which we purport to result from a corresponding variance in the timing of inhibitory neural processing after the stop-signal). Indeed, while the stop-signal locked bin-quantification in our study also didn’t show significant increases in burst rates between stop and go trials, quantifying β bursts in each individual’s SSD-to-SSRT period revealed clear differences.

In addition to this early-latency β bursting in the SSD-SSRT period, we notably also observed clear increases in β bursts at post-SSRT latencies in both the STN and thalamus. While we had no hypothesis about such a finding a priori, we surmise that these later peaks in β bursting may relate to the slower activation of the basal ganglia indirect pathway during movement cancellation (Jahfari et al., 2011; Sano et al., 2013; Schmidt et al., 2013; Mallet et al., 2016). Indeed, one recent framework of movement cancellation, supported by a body of neurophysiological work in rodents, contains the proposition that stopping is a two-step and not a unitary process (Schmidt et al., 2013; Schmidt and Berke, 2017). This two-step model, termed the “Pause then Cancel” model, consists of two phases. An initial Pause involves rapid gating of STN through hyperdirect pathway activation. Meanwhile, a parallel Cancel phase implements indirect pathway activation to eliminate drive to movement from the direct pathway. The STN plays a critical role during both of these phases. Indeed, computational modeling of the human basal ganglia has suggested that inhibitory control could rely on activation of both the hyperdirect and indirect basal ganglia pathways in parallel. While the former implements the rapid gating of the STN, thereby raising the response threshold and buying time for the evaluation of conflicting motor programs, the latter ultimately removes drive to movement from the direct basal ganglia pathway (Frank, 2006; Wiecki and Frank, 2013). If β bursts do in fact relate to both hyperdirect and indirect pathway activation, this has major implications for emerging theories of movement cancellation. Within the current dataset, the subcortical β burst dynamics are in line with this proposition, making them a candidate signature for a unitary inhibitory signal that coordinates both hyperdirect and indirect pathway inhibition.

### STN β bursts influence motor output by raising SMC burst rates

By leveraging unique simultaneous recordings of SMC and subcortical regions, we found that early bursts in STN following stop signals influence motor output by increasing β burst rates in SMC, moreover within around 50ms. Conversely, there was no increased likelihood of STN bursting following SMC bursts, suggesting a degree of directionality that is in line with the proposition that signaling from STN ultimately causes an upregulation of inhibition at the level of SMC. Notably, and somewhat surprisingly, the same was not the case for thalamic β bursts, which were not consistently followed by SMC bursts. This finding suggests that β bursts in STN directly influence β burst rates in SMC. Given that β bursts in SMC have an inhibitory influence on motor output (Little et al., 2019), this points to a possible mechanism by which the STN can influence motor output during movement cancellation.

Previously, we reported that averaged β power in the STN has a Granger-causal relationship to β power in SMC during motor conflict. Moreover, the degree to which STN β predicts SMC β correlates with behavioral measures of the level of motor conflict (Wessel, Waller, and Greenlee, 2019). Results from the current study demonstrating increased SMC β bursting following cancellation-associated STN bursts are in line with this assertion that the STN has direct influences on SMC that relate to the degree of inhibitory control being deployed.

### STN β bursts precede thalamic bursts during movement cancellation

In line with the proposed cortico-subcortical cascade underlying inhibitory control, according to which cortical signals to STN lead to inhibition of the thalamus via the GPi, we found that β bursts following the stop signal across our study sample occur in STN earlier on average than in the thalamus. Despite the fact that the VIM group showed a faster cancellation process (indexed by SSRT) compared to the STN group, the first β burst after the stop-signal occurred later on average in that group. A subsequent assessment of subcortical burst latency in the patient with both STN and VIM recordings confirmed that stop signal-related STN β bursts on average occur prior to VIM bursts within the same individual. Notably, this finding was a result of a highly unusual case, wherein VIM implants were removed to place STN implants, lending an opportunity to record from both regions at once. This evidence lends insight into inhibitory control circuits that would be impossible to obtain under any other circumstances or by using noninvasive methodological approaches in humans.

Taken together, our results suggest that thalamic β bursting is first observed following initial bursts in the STN, and that bursts in the STN and thalamus both relate to inhibitory processing, which precedes SSRT. In addition, STN bursting leads to rapid upregulation of bursts in SMC, whereby inhibitory effects on motor output may be realized. It was surprising that bursts in the thalamus did not have a similar relationship with SMC β bursts, especially given the apparent association of bursts in the thalamus with inhibitory control at early latencies. We note that it is possible that our definition of time bins could potentially segment the data in such a way that these relationships were not observed. It is also possible that the STN has separate, parallel inhibitory effects on SMC and thalamus, so that the rapid gating of SMC by STN does not require signaling *through* the thalamus. In this way, the STN could “buy time” for movement cancellation by inhibiting SMC, while thalamic inhibition by the STN removes the drive to movement.

This interpretation would align closely with a two-stage, Pause-then-Cancel model of inhibition, derived based on neurophysiological recordings of basal ganglia nuclei in rodents (Schmidt and Berke, 2017; Schmidt et al., 2013).

### Broad and clinical implications

Here, we present evidence of an association between β bursts and movement cancellation in all recorded regions of a fronto-basal ganglia pathway for inhibitory control, further establishing transient β as a candidate signature of inhibitory control in the brain. β bursts carry fine-grained information about motor output on the single-trial level, making this signature a powerful tool for investigating the nature and timing of control in the brain. They relate more closely to underlying mechanisms at the level of the cortex than do average β changes (Sherman et al., 2016), and therefore allow researchers who study movement and control to relate cognitive and motor processes not only to changes at the systems level, but to specific changes at the level of neuronal populations.

These findings also hold particular relevance for emerging clinical treatments of movement disorders. Adaptive deep brain stimulation (aDBS) is an intervention being tested for the treatment of motor symptoms in movement disorders such as Parkinson’s disease. This type of DBS is designed to suppress abnormally high levels of β power in STN that relate to motor symptoms by stimulating only when sensors are activated by long, high amplitude β bursts. This approach is advantageous compared to continuous DBS because it reduces potential unwanted side effects through reduced stimulation, lengthens the window of therapeutic effect, and maximizes DBS battery power over the lifetime of the device (Little et al., 2013; Little et al., 2016; Meidahl et al., 2017). Though recent testing of aDBS suggests that these stimulation protocols work by curtailing pathological and not physiological bursts – and may in fact spare more physiological β than continuous DBS (Tinkhauser et al, 2017) – no studies have investigated the effects of aDBS on inhibitory control processes. More research of aDBS effects on physiological bursts during inhibitory control would rule out potential complications for control processes. Moreover, such research would provide the opportunity to evaluate the effects of physiological and pathological β bursts in the STN on network-wide β bursts and behavior.

### Limitations

We note several limitations to the current work. Firstly, the use of a movement disorder patient population was necessary to obtain deep brain recordings, but consequentially these findings may not generalize entirely to healthy individuals. Our participants were undergoing brain surgery at the time they performed the behavioral task, which may have caused distraction or fatigue. At the beginning of the surgery, patients were administered dexmedetomidine, an intravenous sedative. Though patients were required to be awake and responding to instructions from the clinical team at least 30 minutes before our recordings and off all sedatives, it is not entirely possible to rule out lingering effects of sedation. These limitations are endemic to the endeavor of human intracranial neurophysiology. Another limitation is the possibility that some β bursts in our data may be non-representative due to the higher number of pathologically long, high-amplitude β bursts identified in patients with movement disorders, especially PD (Tinkhauser et al., 2018; Lofredi et al., 2019). However, the use of a high cut-off amplitude threshold to identify β bursts (such as the one we used here) has been shown to lead to selection of a shorter subset of β bursts (Schmidt et al., 2020), maximizing our chances of selecting non-pathological bursts from the data.

### Conclusion

In conclusion, this study provides the first network-level neurophysiological evidence for a proposed cascade of cortico-subcortical processing according to which β band burst-like signals between basal ganglia, thalamus, and SMC are underpinning the human ability to rapidly stop an action. This was achieved using a highly unique sample of multi-site intracranial recordings – including a simultaneous recording from both subcortical sites – that is unprecedented in human cognitive neuroscience studies. We found that both STN and thalamus showed increased β burst signaling in the critical time period following the stop-signal, and that STN bursts in particular were followed at low latency by β bursting in SMC. Given that these SMC bursts have been associated with an inhibited state of the motor system (Little et al., 2019; Soh et al., 2021) this strongly speaks in favor of the theory that action stopping is achieved via a rapid re-instantiation of inhibitory control following β burst signaling from the subcortical basal ganglia. In addition, β bursts in STN temporally preceded bursts in thalamus, supporting circuit models of inhibitory control which propose that inhibitory STN activity precedes activity in the thalamus. These findings further confirm transient β bursts as a signature of inhibitory control in the human brain.

## Acknowledgements

The authors would like to thank Haiming Chen for his assistance with surgical recordings, as well as the patients in this study for volunteering their time. This research was funded by an NIH fellowship (T32GSMC08540) to DAD, Carver College of Medicine/Iowa Neuroscience Institute Research Program of Excellence funding to JDWG and JRW, and grants from the NIH (R01NS117753) and NSF (CAREER 1752355) to JRW.

## Author contributions

DAD conducted the experiments, JDWG conducted surgical aspects of the experiments (i.e., placing recording electrodes), JRW and JDWG designed the experiments, and DAD and JRW wrote the paper.

## Method

### Participants

Twenty-three adult patient research participants were recruited at the University of Iowa Hospitals and Clinics over a two-year period. Participants were recruited from all neurosurgical candidates slated for DBS electrode implantation in the thalamus (specifically, the ventral intermediate medial nucleus, VIM) or STN. All patients sought DBS treatment for a movement disorder. STN DBS patients typically had a diagnosis of idiopathic Parkinson’s disease and VIM DBS patients typically had a diagnosis of essential tremor. One patient, with diagnoses of both essential tremor and Parkinson’s disease, had existing VIM implants removed and STN implants placed within the same surgery. From this patient, we recorded data from unilateral VIM and bilateral STN.

Two participants’ data were excluded from analyses based on behavioral performance (more information about exclusion criteria can be found in the section titled “Behavioral analysis”), leaving a sample of twenty-one participants (9 female, mean age: 67 years, age range: 52-78). Information on participant handedness, laterality of symptoms, and motor symptom severity scores for participants included in analyses can be found in **Supplementary Table 1**. When available, Unified Parkinson’s disease rating scale (UPDRS; Movement Disorder Society Task Force, 2003) part III motor examination total scores are included for PD patient participants and Fahn-Tolosa tremor scale scores (Fahn, Tolosa, and Marin, 1993) are included for essential tremor patient participants. UPDRS scores are presented as totals of 33 scored items with possible scores of 0-4 per item, and Fahn-Tolosa scores are presented as a total of 21 scored items, with a possible score of 0-4 for each. These experimental protocols were approved by the University of Iowa’s Institutional Review Board (#201402720).

### Data collection procedure

Participants signed a written informed consent document during a clinic visit prior to surgery. Participants knew that research activities were non-essential to their clinical care and that they were free to discontinue research participation at any time, for any reason. Data collection for this study took place during awake bilateral DBS lead implantation surgery. Before surgery began, participants practiced the behavioral task until they were able to perform the task accurately and confidently enough to demonstrate understanding of the instructions. During surgery, two recording sessions took place. Following placement of bilateral subgaleal 4-contact electrode strips (Ad-Tech, Inc) directed posteriorly from the burr holes at the coronal suture so as to sit over SMC, a short recording session (a functional localizer) was used to confirm correct placement of the strip electrodes. Participants performed a short, 40-trial version of the stop-signal task that did not include any stop-signals (i.e., it was purely a two-alternative forced-choice reaction time task). These data were analyzed immediately in the operating room and the electrode lead placement was changed if the initial placement did not reveal the typical signature of SMC activity during movement execution (criteria for optimal placement and localization analysis is described further in the section titled “Analyzing local field potentials”). Then, after the DBS leads (3387, Medtronic, Inc, Minneapolis, MN) were successfully implanted into the bilateral subcortical sites (STN or VIM) using framed indirect stereotactic targeting refined by standard confirmatory physiologic testing (i.e. microelectrode recording, macrostimulation) to define the borders of each nucleus, a second recording session took place, which contained the main experiment (see next section).

### Behavioral paradigm

Participants completed an auditory stop-signal task in the operating room while recordings were collected. Task stimuli were played through in-ear headphones (ER4 SR model with ER38-14F foam buds, Etymotic Research, Elk Grove Village, IL, USA) connected to a Dell laptop running Fedora, using the PsychToolbox package (version 3; Brainard, 1997) in MATLAB (MathWorks, Natick, MA). Participants responded using two large USB response buttons held in the hands (Kinesis Savant Elite 2, Kinesis, Bothell, WA). Participants heard a tone cuing a response (the go signal) every four seconds. This go signal was a 500Hz sine wave tone lasting 100ms. Half of the go signals were presented in the left ear and half in the right ear (in random order); participants were instructed to respond with the button that indicated the side to which the tone was presented. (If the tone was presented in the left ear, the participants pressed the left button, and vice versa.) Participants had 2 seconds to respond to the tone – any response after this deadline was not registered and the task proceeded to a 2s inter-trial interval.

On one-third of trials, participants heard a second, higher pitched tone presented in both ears following the go signal – this was the stop signal, which indicated participants needed to try to cancel their initiated response. The stop signal was a 1500Hz sine wave tone lasting 100ms.

The delay between the go and stop signals, the stop-signal delay (SSD), was adjusted throughout the task to ideally converge on a stopping accuracy of 50%. To this end, the initial SSD was set to 250ms and adjusted in 50ms increments for each hand – failed stop trials resulted in 50ms being subtracted from the SSD and successful stop trials resulted in an addition of 50ms to the SSD. In an attempt to prevent proactive strategies (where participants waited to see if a stop-signal would occur before responding), they were instructed that it was equally important to 1) respond as fast as possible, and 2) try to cancel movements successfully when the stop-signal occurred. The pre-surgical practice with the experiment consisted of one block of 30 trials (10 stop). The main task and recording block following macroelectrode lead placement included four blocks of 48 trials (16 stop). Between each block of the main task, the participants rested for several seconds as needed and received feedback on their performance if necessary. For a diagram of the auditory stop-signal task, see ***Figure 1***.

### Local field potential recordings

Local field potentials (LFPs) were recorded from the VIM or STN using four macroelectrode contacts on each DBS lead and from two four-, six-, or eight-contact strip electrodes placed in the subgaleal space over SMC (Ad-Tech, Oak Creek, WI; 10 mm spacing center-to-center, 3 mm exposed contact diameter). Most SMC recordings were obtained with four-contract strip electrodes. Information about the number of contacts in subgaleal strips for each participant and the pair of electrodes ultimately chosen for analysis can be found in **Supplementary Table 2**. Based on head box and amplifier limitation on number of channels that can be recorded, we only recorded from two subgaleal channels per hemisphere for the patient with both STN and VIM leads.

The neurosurgeon (JDWG) inserted the motor strips into the subgaleal space posterior to the stereotactic burr hole at the coronal suture, para-sagitally in direction and anterior-posterior in alignment to cover the precentral gyrus. Estimations of the most posterior electrode were ∼6 cm posterior to the coronal suture, which is consistent with a posterior placement covering precentral gyrus and SMC (Park et al., 2007; Rivet et al., 2004). We used the same electrode placement procedure for an identical recording set-up described in Wessel, Waller, and Greenlee (2019). LFP recordings were made on a Tucker-Davis technologies (Alachua, FL) system, using a RA16PA 16-Channel Medusa pre-amplifier and a RA16LI head-stage. The sampling rate for recording was 24Hz or 2Hz, with a low-pass filter of 7.5 kHz on the hardware side. Stimulus onsets were marked in the recording using a TTL pulse from a USB Data Acquisition Device (USB-1208FS, Measurement Computing, Norton, MA) triggered by the stimulus presentation laptop.

### Preprocessing local field potentials

Preprocessing and analysis of LFP data was conducted using custom MATLAB scripts. Data, task code, and analysis scripts for this study can be found on the Open Science Framework at [*link will be added at time of publication*]. Electrical line noise from the operating room environment was filtered from the data using EEGLAB’s (Delorme and Makeig, 2004) *cleanline()* function after which the recordings were down-sampled to 1000Hz for analysis. Then, the recordings were visually inspected for any artifacts. Any 1s segment of the recording containing an artifact was marked for removal and thrown out of the data prior to subsequent analyses.

### Analyzing local field potentials

For all analyses, electrode pairs were converted to bipolar montages, resulting in three bipolar recordings from each side of the subcortical location (channel 0 – channel 1, channel 1 – channel 2, channel 2 – channel 3). We conducted LFP analyses specifically using the bipolar array located in the VIM proper (determined by clinical testing) and the bipolar array in the STN which included the greatest number of β bursts over the entire recording. We utilized the Medtronic lead labeling nomenclature such that contact 0 was the distal-most contact and positioned at the ventral border of each nucleus, contact 3 was the most proximal contact. Intercontact spacing was 1.5 mm. For thalamic DBS leads, placement of contacts 0 and 1 in the VIM were confirmed with clinical stimulation and testing in the operating room. However, based on the trajectory of implantation, contacts 2 and 3 were still located in the motor thalamus, albeit a more dorsal subsection (likely the ventral oral posterior region of the ventral lateral nucleus). Accordingly, contacts 2 and 3 were analyzed in a similar manner to the VIM contacts (0 and 1) and these results are included in the Supplementary text.

Broadband event-related spectral perturbation (ERSP) plots of go-locked activity were made using a window of 100ms before stimulus onset to 1500ms following stimulus onset. A baseline window of 500ms to 200ms before stimulus onset was used to perform baseline corrections. Data were converted to time-frequency series using the filter-hilbert method: a Hilbert transform was applied to data filtered at specific frequencies (1-50Hz) with a window of .5Hz below and above that frequency using symmetric 2-way least-squares finite impulse response filters. The analytic signal was extracted by computing the squared absolute value of the complex signal.

This ERSP analysis procedure was also used to check subgaleal strip placement in the operating room. Following localization, go-signal-locked ERSP plots were created for each bipolar array on the two subgaleal strips and visually inspected. One of the researchers in the OR (DAD) visually checked these ERSP plots for a visible, circumscribed decrease in average β band amplitude, a signature of movement-related activity in SMC (Pfurtscheller and Lopes da Silva, 1999). If no β suppression was observed in any bipolar array on one or both strips, those strip electrodes were replaced by the neurosurgeon (JDWG) for more optimal positioning. Repositioning of one strip electrode was required in two of the 21 participants and repositioning of both was required in one participant. Time constraints did not allow for a repeat of the localizer task after the strips were replaced. However, examination of main task recording data from participants who had strips replaced showed β suppression in subgaleal contacts during responding on go trials (**see Figure 2**), suggesting that repositioning did result in improved signal from SMC.

### B burst quantification

To classify β bursts, detection was performed using the same procedure as in Wessel (2020) and Shin et al. (2017). Data from each bipolar electrode array were convolved with a complex Morlet wavelet constructed using the following equation:

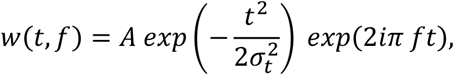

With 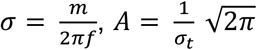, and m = 7 (cycles) for each frequency in the β band (15–29 Hz). The absolute value of the resulting complex data was squared to yield time-frequency power estimates. The resulting time-frequency data were epoched around events of interest (go and stop signals) with a window of 500ms before stimulus onset to 1000ms after stimulus onset. B bursts were classified by identifying local maxima in the trial-by-trial time-frequency data that exceeded six times the median of the time-frequency power for that specific electrode across the recording and that lasted at least two β cycles.

### Statistical analysis

#### Behavioral analysis

Two participants’ data were excluded from analysis because they performed below chance accuracy (50%) on go trials or did not perform the task correctly during the main task (i.e., did not stop successfully to a single stop signal during more than one block). Participants included in the final analysis therefore included 9 STN DBS patients, 11 VIM DBS patients, and one participant with both VIM and STN DBS. With the exception of the two key analyses that made use of the simultaneous recording of STN and VIM (i.e., quantifying bursts in subcortical regions between SSD and SSRT, and comparing latencies of bursts in STN and VIM in the single-subject analysis), the patient with both regions recorded was only included in one of the sample groups – in other words, the participant with data from both STN and VIM only contributed data to the STN group for most statistical comparisons. Individual task blocks within participants were excluded from analysis if mean accuracy on go trials during a block was less than 60%, or if participants did not successfully stop on at least one stop trial. Based on these criteria, six of the 21 participants had one block of four excluded from behavioral and LFP analysis. The remaining fifteen subjects retained all of their data. Mean accuracies for stop and go trials were extracted for each subject. Go trials were considered incorrect if participants pressed the wrong button or missed responding before the 2s deadline. Mean RTs for failed stop and successful go trials were extracted for each subject, and SSRT was calculated using the integration method with go omission replacement (Verbruggen et al., 2019). Differences in average accuracy and RT measures between STN and thalamic implant patient groups were tested using two-sample *t*-tests.

#### Temporal progression of β Bursts

In all analyses of the temporal progression of β bursts, we included data from subcortical sites both ipsi- and contralateral to the correct trial response. This approach is supported by findings that STN has a bilateral representation during movement execution (Alegre et al., 2005; Devos et al., 2006). Data from SMC was only included from sites contralateral to the correct trial response (except for the contralateral compared to ipsilateral hemisphere analysis in **Figure 3**). To quantify progression of β bursts across time following go- and stop-signals, counts of bursts across all trials of the same type (correct-go, failed stop, or successful stop) were binned by burst latency with respect to stimulus onset latency, in bins of 100ms from stimulus onset to 900ms following stimulus onset. For stop trials, latencies of β bursts were calculated as the latency difference between burst time and time of stop signal onset. For matched go trials, β burst latency was calculated in respect to when the stop-signal would have occurred if the trial was a stop trial (i.e., the SSD set in the staircase for that trial). In addition to quantifying burst rates following the stop signal, we quantified bursts during a pre-stop signal baseline to ensure that there were no differences between burst rates across conditions before the stop-signal was presented. To quantify pre-stop baseline bursts, we summed bursts in the 100ms before SSD and averaged by the total number of trials. Trials with a SSD of 50ms were excluded from this baseline quantification because there were not 100ms of time between the go signal and stop signal to serve as a pre-stop baseline.

For the analysis in which bursts were time-locked to β bursts at another recording site, the analysis was constrained to bursts within 500ms following the stop-signal (or 500ms following SSD for matched-go trials) in order to assess bursts that would reasonably contribute to movement cancellation based on average sample SSRT (474ms; **Table 1**). For the *first* subcortical β burst within 500ms following the stop signal, we calculated the latency difference between the subcortical burst and all bursts in SMC during the same trial. This analysis was also repeated with the reverse “directionality”, analyzing subcortical burst rates time-locked to SMC β bursts in the same manner.

Permutation-based statistics were used to evaluate statistical significance. Specifically, two-way repeated measures ANOVAs were calculated with factors of *TIMEPOINT* and *TRIAL TYPE*. ANOVAs were bootstrapped by comparing the resulting *F* values for each factor to null distributions of *F* values from 10,000 tests with data labels randomized. True *F* values were considered significant if they were greater than the *F* value at the 95^th^ percentile of the null distribution and *p*<.05. Pairwise differences between successful stop and go trials and successful and failed stop trials were calculated using *t*-tests with Bonferroni-Holm corrections for multiple comparisons.

#### Specificity of evoked burst rate changes to selected motor contacts

The pair of two adjacent subgaleal electrodes containing the greatest number of β bursts across all subgaleal contacts were selected for analysis. To check whether the burst rate changes observed in our main analysis were specific to the selected subgaleal electrodes (and therefore indicative of specificity of these changes to certain parts of SMC), we plotted go-signal and stop-signal locked burst rates for selected subgaleal channels and the adjacent pairs of electrodes (when selected channels were not the first or last in the array). Selected channels for both hemispheres on average had the greatest number of β bursts following the go and stop signal, displayed rapid decreases in β bursts during movement, and reflected the same early increase in cancellation-related bursts as in STN. During movement execution and cancellation, the selected channels accounted for the fastest rate of change in early time bins in the right but not left hemisphere. These results are presented in the **Supplementary Figures 1 and 2**.

#### Comparing reaction times for trials with and without β Bursts

In order to assess the effect of β burst presence in SMC on reaction time for trials containing a response (correct go and failed stop trials), we compared the average reaction times for trials that contained no bursts or at least one burst in SMC. Trials that contained no bursts between go signal onset and reaction time were considered “no burst” trials, while trials containing at least one burst between the go signal and the response were considered “burst trials”. A bootstrapped, two-way repeated measures ANOVA was conducted with factors of *TRIAL TYPE, BURST PRESENCE*, and *TRIAL TYPE × BURST* interaction.

#### Calculating bursts in SSD-SSRT period

To calculate subcortical burst rate differences between failed and successful stops and matched go trials in the SSD-SSRT delay, we quantified the total number of bursts for each trial type between trial-specific SSD and the participant’s average SSRT. The participant with both VIM and STN recordings contributed both STN and VIM recording data to this analysis. We conducted a bootstrapped ANOVA with *TRIAL TYPE* as a factor. Follow-up pairwise comparisons between successful and failed stops were conducted using *t*-tests with Bonferroni-Holm corrections for multiple comparisons.

#### Timing of β bursts at each subcortical recording site

To assess the relative timing of bursts at different sites during stop trials, we calculated the average latency of first bursts with respect to subject SSRT at each recording location following stop-signal onset. Though we present STN and VIM timing together in the same figure for sample averages (**Figure 6**), note that all but one participant had either STN or VIM recordings, not both. This analysis was also performed in the subject with STN and VIM recordings – these results are presented separately in the second panel of **Figure 6**. We also conducted a between- and within-subjects ANOVA with a within-subject factor of *TRIAL TYPE* and between-subjects factor of *SUBCORTICAL LOCATION* to assess whether successful stops might be associated with a shorter delay between STN and VIM bursts than failed stops (again, acknowledging that these regions were recorded in separate individuals).

## Supplemental information

**Supplementary Table 1.** Information about participant diagnoses, handedness, symptom laterality, and pre-operative symptom severity scores. For participants with Parkinson’s disease, scores shown are from the motor examination portion (i.e. total part III) of the Unified Parkinson’s disease rating scale (UPDRS). For essential tremor participants, scores are from the Fahn-Tolosa tremor scale.

| Participant number | Diagnosis | Implantation site | Handedness | Symptom laterality | Symptom severity score |
| --- | --- | --- | --- | --- | --- |
| 1 | Parkinson's disease | STN | R | - | - |
| 2 | both | STN/VIM | L | left | 49 |
| 3 | Parkinson's disease | STN | R | bilateral | 38 |
| 4 | Essential tremor | VIM | R | - | - |
| 5 | Parkinson's disease | STN | R | left | 33 |
| 6 | Essential tremor | VIM | R | right | 47 |
| 7 | Essential tremor | VIM | R | right | 59 |
| 8 | Parkinson's disease | STN | R | right | 37 |
| 9 | Essential tremor | VIM | R | left | 43 |
| 10 | Essential tremor | VIM | R | right | - |
| 11 | Parkinson's disease | STN | R | right | 34 |
| 12 | Parkinson's disease | STN | R | right | - |
| 13 | Essential tremor | VIM | R | bilateral | 78 |
| 14 | Essential tremor | VIM | R | right | 85 |
| 15 | Essential tremor | VIM | R | left | - |
| 16 | Parkinson's disease | STN | R | right | 31 |
| 17 | Essential tremor | VIM | R | bilateral | 69 |
| 18 | Parkinson's disease | STN | R | right | - |
| 19 | Parkinson's disease | STN | R | left | 29 |
| 20 | Essential tremor | VIM | R | right | 66 |
| 21 | Essential tremor | VIM | R | bilateral | 57 |

**Supplementary Table 2.** Information about number of subgaleal contacts used for each participant and pairs of contacts selected for main analyses.

| Participant number | Number of subgaleal contacts per side | Pairs selected for analysis, (left/right) |
| --- | --- | --- |
| 1 | 4 | [1, 2] / [1, 2] |
| 2 | 2 | [2, 3] / [2, 3] (only channels recorded) |
| 3 | 4 | [1, 2] / [1, 2] |
| 4 | 4 | [1, 2] / [2, 3] |
| 5 | 4 | [1, 2] / [3, 4] |
| 6 | 4 | [3, 4] / [2, 3] |
| 7 | 4 | [1, 2] / [1, 2] |
| 8 | 6 | [1, 2] / [5, 6] |
| 9 | 6 | [5, 6] / [5, 6] |
| 10 | 8 | [7, 8] / [1, 2] |
| 11 | 4 | [2, 3] / [3, 4] |
| 12 | 4 | [2, 3] / [3, 4] |
| 13 | 4 | [3, 4] / [3, 4] |
| 14 | 4 | [1, 2] / [1, 2] |
| 15 | 4 | [2, 3] / [1, 2] |
| 16 | 4 | [1, 2] / [3, 4] |
| 17 | 4 | [3, 4] / [1, 2] |
| 18 | 4 | [3, 4] / [1, 2] |
| 19 | 4 | [3, 4] / [1, 2] |
| 20 | 4 | [1, 2] / [2, 3] |
| 21 | 4 | [3, 4] / [3, 4] |

**Supplementary Figure 1.**
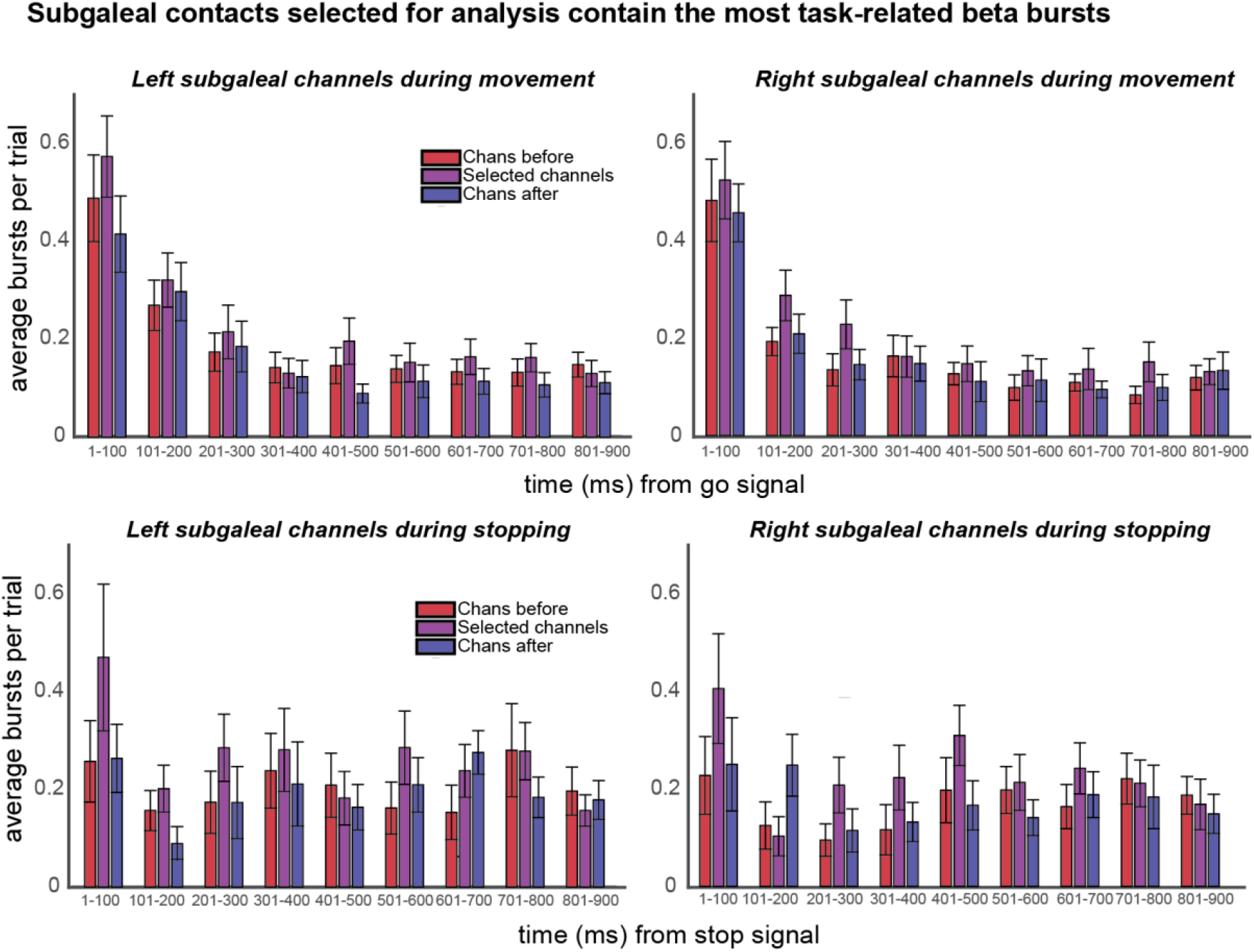
Beta burst rates for all subgaleal electrode pairs following the go signal during go trials and following the stop signal during stop trials.

**Supplementary Figure 2.**
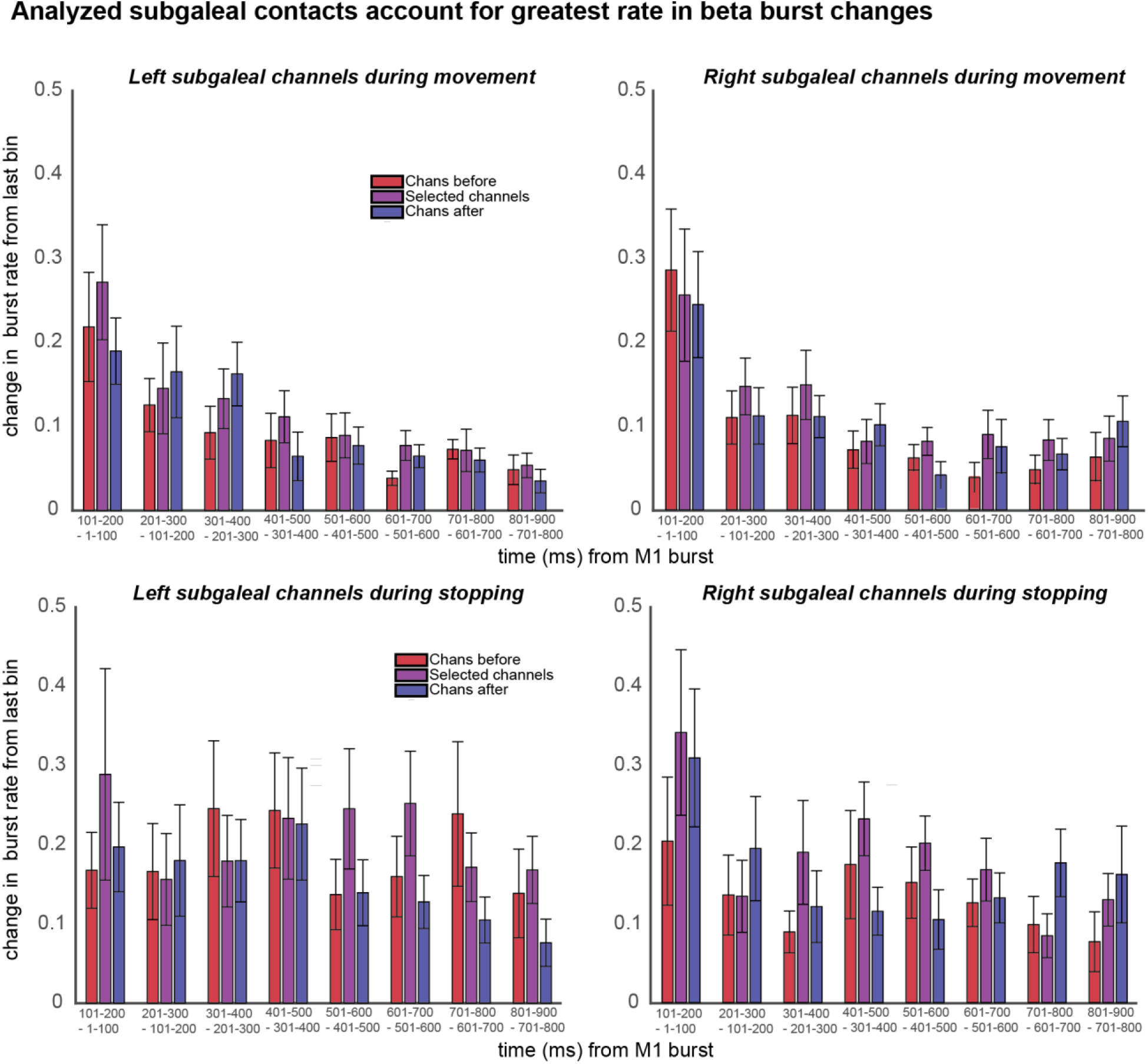
Rate of change in beta bursts during going and stopping in all subgaleal contact pairs. In the right hemisphere, selected channels account for the greatest rate of change in beta bursts. In the left hemisphere, selected channels on average do not account for the greatest rate in change, though differences between rate of change in selected and non-selected channels were not significantly different.

**Supplementary Table 3.** Electrode channels on subcortical DBS leads that contained the highest number of beta bursts during the recording session. For STN participants, these channels were used for beta burst quantification. For VIM participants, contacts 0 and 1 were used based on anatomical consideration.

| Participant number | Implantation site | Channels selected in left hemisphere | Channels selected in right hemisphere |
| --- | --- | --- | --- |
| 1 | STN | 1, 2 | 1, 2 |
| 2 | STN/VIM | 1, 2 | 2, 3 |
| 3 | STN | 1, 2 | 1, 2 |
| 4 | VIM | 1, 2 | 1, 2 |
| 5 | STN | 1, 2 | 2, 3 |
| 6 | VIM | 2, 3 | 1, 2 |
| 7 | VIM | 0, 1 | 0, 1 |
| 8 | STN | 1, 2 | 2, 3 |
| 9 | VIM | 0, 1 | 0, 1 |
| 10 | VIM | 1, 2 | 2, 3 |
| 11 | STN | 0, 1 | 0, 1 |
| 12 | STN | 0, 1 | 1, 2 |
| 13 | VIM | 0, 1 | 0, 1 |
| 14 | VIM | 0, 1 | 1, 2 |
| 15 | VIM | 0, 1 | 0, 1 |
| 16 | STN | 1, 2 | 1, 2 |
| 17 | VIM | 0, 1 | 2, 3 |
| 18 | STN | 1, 2 | 2, 3 |
| 19 | STN | 2, 3 | 0, 1 |
| 20 | VIM | 0, 1 | 0, 1 |
| 21 | VIM | 2, 3 | 0, 1 |

**Supplementary Figure 3.**
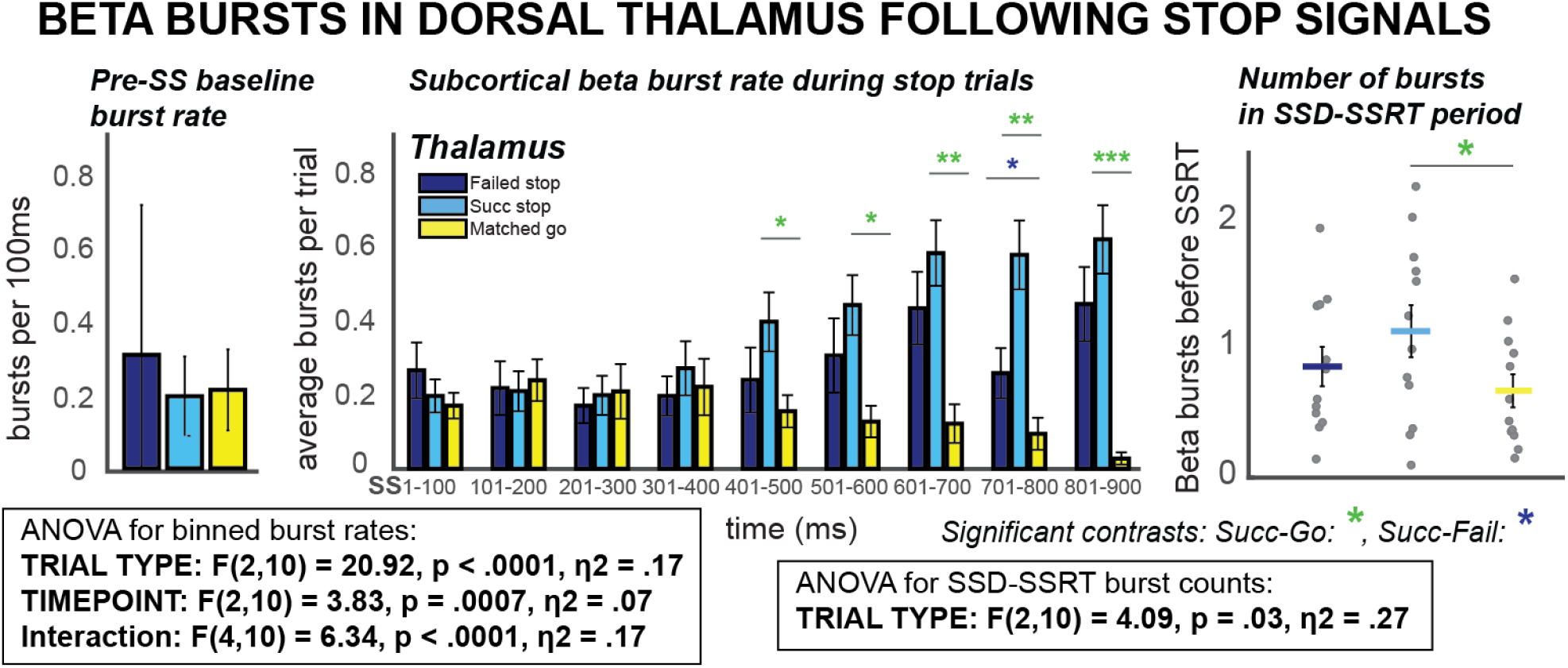
Beta burst rate for all trial types following the stop-signal in dorsal thalamus. Beta bursts increase slightly at early latencies, so that successful stop trials contain more beta bursts than matched go trials before SSRT. Later peaks in beta bursting are also observed.

**Supplementary Figure 4.**
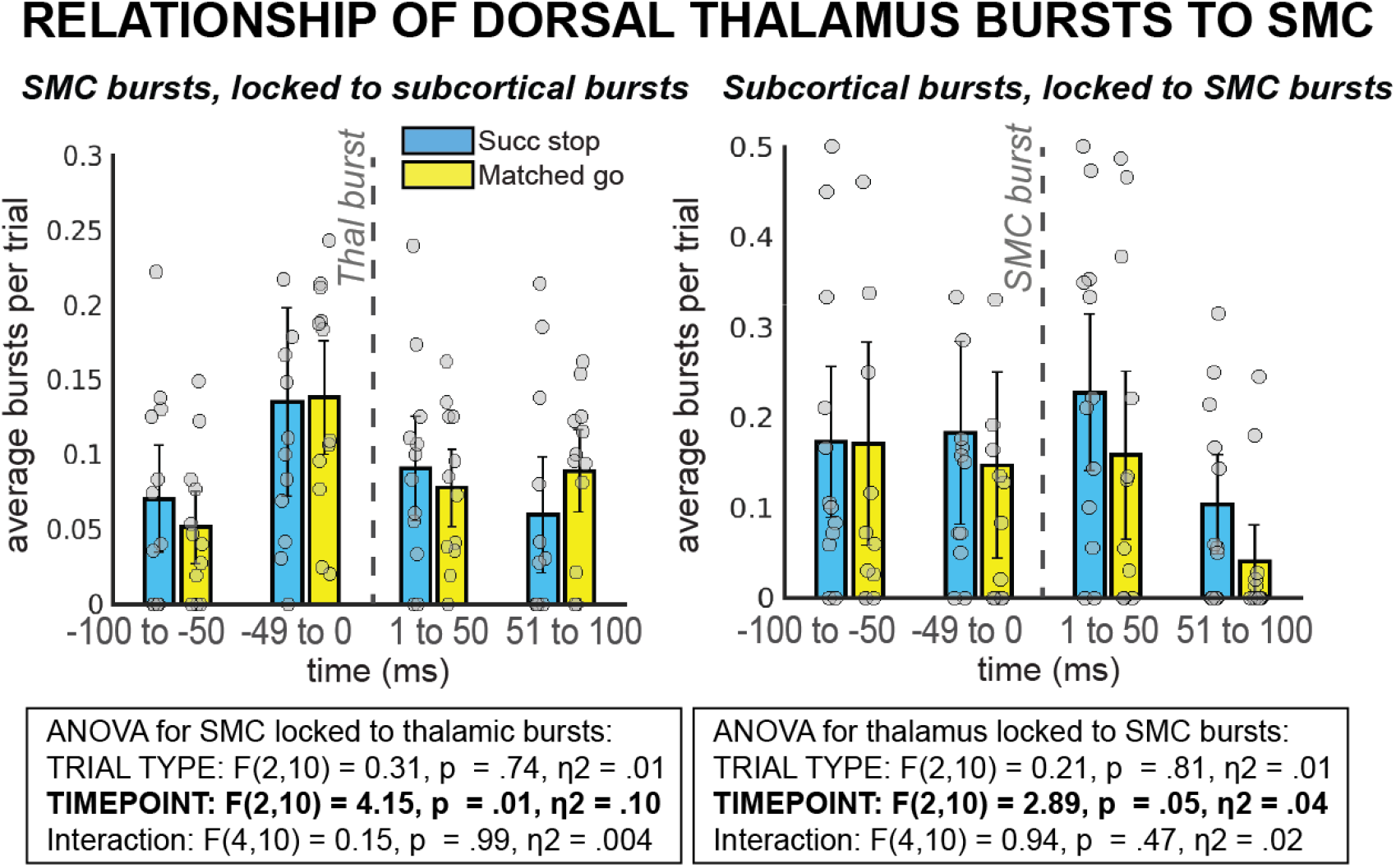
Beta burst rates time-locked to preceding beta bursts in dorsal thalamus and sensorimotor cortex. There is not a time-resolved temporal relationship between bursting in dorsal thalamus and sensorimotor cortex.

